# Genetic Control of Prickles in Tetraploid Blackberry

**DOI:** 10.1101/2024.11.27.625708

**Authors:** Carmen A. Johns, Alexander Silva, T. Mason Chizk, Lacy Nelson, John R. Clark, Rishi Aryal, Hudson Ashrafi, Ellen Thompson, Michael Hardigan, Margaret Worthington

**Affiliations:** University of Arkansas System Division of Agriculture, Department of Horticulture, 316 Plant Sciences Building, Fayetteville, AR 72701; North Carolina State University, Department of Horticulture, 2721 Founders Dr, Raleigh, NC 27607; Hortifrut Genetics, LTD. 589 Casserly Road, Watsonville, CA 95076; United States Department of Agriculture, Agriculture Research Service, Horticultural Crops Production and Genetic Improvement Research Unit, 3420 NW Orchard Avenue, Corvallis, OR 97330

**Author notes:** Corresponding author: Margaret L. Worthington.

**Keywords:** *Rubus* subgenus *Rubus*, thorns, GWAS, KASP, marker-assisted selection, autopolyploid, bramble, caneberry

## Abstract

Prickle-free blackberry (*Rubus* subgenus *Rubus*) canes are strongly preferred by growers due to food and worker safety concerns and damage to fruit from mechanical injury by prickles. This project was conducted to identify the genetic region responsible for prickle-free canes derived from the recessive ’Merton Thornless’ source in autotetraploid blackberry using a genome-wide association study, develop diagnostic KASP markers for prickle-free canes, and determine the effects of allele dosage at the prickle-free locus on prickle density in two biparental populations. The prickle locus was located on chromosome Ra04 from 30.48 to 36.04 Mb in an extensive LD block, with the peak SNP located at 33.64 Mb. Two diagnostic KASP markers were developed that correctly predicted the phenotype of 97% and 96% of 626 diverse fresh-market blackberry genotypes from multiple breeding programs, respectively. Allele dosage at the prickle-free locus had a significant impact on prickle density, with duplex prickly genotypes having significantly higher prickle density than simplex genotypes in both biparental populations studied. Five potential candidate genes with functional annotations related to epidermal, trichome, and/or prickle development were identified within the prickle-free locus, but no nonsynonymous polymorphism within these genes were identified.

**Article Summary:** Blackberry cultivars with prickle-free canes are strongly preferred by growers and shipper marketers. The objectives of this study were to map the prickle-free locus in tetraploid fresh-market blackberries using a genome-wide association approach and develop diagnostic molecular markers that breeders can use to plan crosses and screen seedlings for prickle-free canes. The prickle-free locus was mapped to a 5.6 Mb region on chromosome Ra04. Two molecular markers developed in this region correctly predicted the phenotype of 96% or more of the breeding selections and cultivars in a large validation panel composed of germplasm from three breeding programs.

## Introduction

Blackberry (*Rubus* subgenus *Rubus*) is a specialty crop with an increasing share of the fresh berry market. The growth in demand and rising production costs have resulted in a need for cultivars that are adaptable to many environments and cultural management approaches. Prickles are a burden in blackberry production systems because they are a food safety concern and can cause mechanical damage to fruit and shorten shelf life (Clark et al. 2007). A common goal of fresh-market blackberry breeding programs is to release cultivars that lack “thorns” or, more correctly, prickles. Often, these terms are used interchangeably in literature. Prickles are botanically differentiated from thorns and spines because they are borne of cortical and epidermal cells (Kellogg et al. 2011). Hereafter, we will use prickles throughout this article.

Prickle-free blackberry cultivars have been developed from multiple sources. Non-heritable sources of the prickle-free trait are borne of a chimera in which the L1 layer of tissue, or epidermis, is responsible for the trait expression. To maintain trueness to type, cells must be propagated only from the mutated apical meristem, rendering this source of the prickle-free trait difficult to use in breeding. There is also a dominant, heritable source of prickle-free blackberries found in the cultivar ’Austin Thornless’. There are major drawbacks to this dominant source, including a strong association with trailing growth habit and canes only being prickle-free over 30 cm in height (Clark et al. 2007).

The most commonly used source of prickle-free canes in fresh market tetraploid blackberries is derived from ’Merton Thornless’. ’Merton Thornless’ was developed at the John Innes Institute in England by crossing a wild prickle-free diploid *Rubus ulmifolius* plant to prickly tetraploid germplasm. While all the initial tetraploid plants developed from the interploidy cross were prickly, crosses among these prickly tetraploids resulted in a small percentage of seedlings with prickle-free canes as expected for recessive inheritance in a multisomic tetraploid (Scott et al. 1957). ’Merton Thornless’ was released from these crosses and used as a heritable, recessive source of prickle-free blackberry canes in many breeding programs, especially in North America (Coyner et al. 2005). The process of developing prickle-free cultivars took many years because introgression of recessive traits is a lengthy process in autopolyploids (Coyner et al. 2005). Furthermore, ’Merton Thornless’ has trailing plant architecture, which conflicted with the goal of erect plant architecture in most fresh-market blackberry breeding programs, adding time to remove the trailing tendency in prickle-free progeny (Clark et al. 2007).

Development and implementation of molecular tools in blackberry breeding programs are needed to shorten breeding cycles and continue growing the industry for this high-demand specialty crop. Erect and semi-erect fresh-market blackberries are autotetraploid, which complicates genetic research (Foster et al. 2019). Molecular breeding studies in autopolyploids are complicated by their high degree of heterozygosity, multisomic inheritance, and challenges with accurately calling allele dosage (Bourke et al. 2018). Recently, multiple software programs that are capable of scrutinizing autopolyploid genetic data have been developed, along with a chromosome-scale reference genome of diploid *Rubus argutus* blackberry (Bourke et al. 2018; Brůna et al. 2023). These advances, along with new genotyping platforms capable of producing thousands of molecular markers with adequate depth of coverage in autopolyploid crops, have made molecular breeding research increasingly practical in tetraploid blackberries (Chizk et al. 2023; Worthington et al. 2024).

The genetic control of prickle-free canes has been explored in previous studies in blackberry and red raspberry (*Rubus idaeus*). Castro et al. (2013) built a linkage map of tetraploid blackberry with 119 simple sequence repeat markers and mapped the prickle-free locus to a large region on the distal portion of LG4, but no broadly diagnostic molecular markers were identified for use in breeding. Most recently, a GWAS conducted in a biparental population of red raspberry placed the prickle-free locus at 33.5 to 35.1 Mb on chromosome 4 (Khadgi and Weber 2021). Promising candidate genes proposed in that region included a trichome birefringence-like 2, a MYB16-like transcription factor, and *AGL30* (a MADS-box protein). All the possible candidate genes identified in the red raspberry prickle locus have a role in epidermal or trichome regulation, development, and/or differentiation (Khadgi and Weber 2020, 2021).

A trichome is an outgrowth that forms from the epidermis of a plant. Trichomes can be categorized into two broad classifications: glandular and non-glandular (Huchelmann et al. 2017). Glandular trichomes form a mass, or “head” that is rich in secondary metabolites (Kellogg et al. 2011; Huchelmann et al. 2017). Kellogg et al. (2011) examined prickle formation on canes of blackberry cultivars ’Arapaho’ and ’Prime-Jim^®^’, red raspberry cultivars ’Heritage’ and ’Canby’, and the rose hybrid cultivar ’Radtko’ (*Rosa hybrida* L.). While blackberry seedlings can have glandular and non-glandular trichomes, only glandular trichomes developed into prickles, and non-glandular trichomes were present on the developing canes and leaves of prickle-free plants (Kellog et al. 2011). Only the plants that had both glandular and non-glandular trichomes present developed prickles. Lignification began at the top of the trichome, working down towards the attachment point on the epidermis (distal-proximal), which indicated that secondary metabolites present in the glandular trichome heads may signal prickle development (Kellogg et al. 2011).

The objectives of this research were to perform a genome-wide association study to more precisely define the locus controlling prickle-free canes derived from ‘Merton Thornless’ in tetraploid blackberries, develop diagnostic KASP markers for the breeding community, and to explore the effect of allele dosage at the prickle-free locus on prickle density in two segregating biparental populations.

## Materials and Methods

### Plant Materials for the Association Study

This study was conducted at the UADA Fruit Research Station (FRS) in Clarksville, AR (lat. 35°31’5”, long. 35°31’5”). Genotypes evaluated for the association study included tetraploid erect and semi-erect blackberry cultivars and UADA breeding selections grown in ten-plant plots. Whole plots were evaluated discretely for the presence or absence of prickles. Two diploid blackberry germplasm accessions [‘Hillquist’ (*R. argutus,* PI 553951) and ‘Burbank Thornless’ (*R. ulmifolius inermis,* PI 554060)] grown in plots at FRS were also included in the prickle GWAS. Ten historical cultivars used as parents in many fresh-market blackberry breeding programs that were not grown at FRS were also included in the GWAS analysis for prickles.

These historical cultivars included ‘Black Satin’, ‘Brazos’, ‘Chester Thornless’, ‘Darrow’, ‘Eldorado’, ‘Loch Ness’, ‘Merton Thornless’, ‘Raven’, ‘Thornfree’, and ‘Triple Crown’. The USDA National Clonal Germplasm Repository provided leaf tissue for these plants and phenotypes were taken from Clark et al. (2007). In total, 374 cultivars and breeding selections were included in the initial association analysis (Supplementary Table 1).

### Genotyping and SNP calling

DNA was extracted from young leaves of each genotype following a modified cetyltrimethylammonium bromide (CTAB) method (Porebski et al. 1997). The samples were quantitated with an Invitrogen Qubit 1 Fluorimeter dsDNA-BR assay kit (Thermo Fisher Scientific, Waltham, MA). Standardized DNA concentrations of 40 µg/mL for each genotype in the panel were submitted to RAPiD Genomics (Gainesville, FL) for Capture-Seq genotyping.

A total of 35,054 custom biotinylated 120-mer Capture-Seq probes were designed using the ‘Hillquist’ (*R. argutus*) reference genome (Brůna et al. 2023). HiSeq2000 was used to perform paired-end sequencing to achieve an average of 5.14 million 150 bp paired-end reads per sample (Chizk et al. 2023). MOSAIK (Lee et al. 2014) was used to align cleaned and trimmed sequencing data to the ‘Hillquist’ genome. Freebayes (Garrison and Marth 2012) was used to perform variant calling. A dataset of biallelic SNP markers with read depths between three and 750 per sample and a minor allele frequency of > 0.01 was generated using VCFtools (Danecek et al. 2011). This dataset was then converted to probabilistic tetraploid allele dosage calls and filtered for quality in UpDog (Gerard et al. 2018). Markers estimated to have a greater than 5% estimated error rate were excluded from the analysis.

### Association Analysis

GWASpoly (Rosyara et al. 2016) was used for association analysis. Binary scores of prickle presence or absence were associated with the filtered set of tetraploid SNPs using a mixed model controlling for population structure with a random effect kinship matrix calculated using the leave-one-chromosome-out (LOCO) method (Listgarten et al. 2012; Yang et al. 2014) and a fixed Q matrix generated by STRUCTURE (Pritchard et al. 2000). The Q matrix was composed of K=6 column vectors determined by the delta K statistic (Evanno et al. 2005) as described by Chizk et al. (2023). Additive and simplex dominance (1-dom) models of gene action were tested. Minor allele frequency and maximum genotypic thresholds were set to 0.05 and 0.95, respectively. LOD significance thresholds of 6.1, 5.3, and 6.1 for additive, simplex alternative allele dominance, and simplex reference allele dominance models were chosen using an α = 0.05 significance threshold and the ‘M.eff’ method. QQ-plots were constructed from association results to compare observed marker scores against nominal P-values.

### Potential Candidate Gene Identification

Patterns of LD decay around the prickle-free locus were analyzed using the *mldest* function in the R package *ldsep* (Gerard 2021) as described by Chizk et al. (2023). A heatmap of the chromosomal linkage disequilibrium (LD) matrix for chromosome Ra04 was constructed using the LDheatmap R package (Shin et al. 2006). Based on the slow rate of LD decay around the prickle-free locus, a 5.6 Mb region extending from the first SNP significantly associated with the prickle-free locus to the distal end of chromosome Ra04 was used to search for potential candidate genes with functional annotations related to epidermal, trichome, and/or prickle development.

### Whole Genome Sequencing

Seventeen fresh-market, tetraploid blackberry cultivars and selections from the UADA breeding program were sequenced with 150 bp paired-end Illumina HiSeq reads at Novogene (Sacramento, CA, USA). This group of sequenced cultivars and selections included four genotypes with prickles and 13 prickle-free genotypes (Supplementary Table 2). Raw reads were cleaned and filtered with Trimmomatic v.0.39 (Bolger et al. 2014) and aligned to the ‘Hillquist’ *R. argutus* genome (Brůna et al. 2023) using BWA-MEM v.0.7.10 (Li 2013). Genetic variant calling was done using GATK4 v.4.2.6.1 using the HaplotypeCaller function (Poplin et al. 2017). 2018). Since fresh-market blackberries are tetraploid, the argument ‘-*ploidy* 4’ was employed to estimate allele dosages based on sequence read counts.

### KASP Marker Development and Validation

Whole genome sequence data was used to identify additional polymorphisms within the genome region significantly associated with prickles in the GWAS analysis. The Hillquist ‘*R. argutus*’ genome was developed using a wild diploid accession with prickles and was assumed to be homozygous for the prickly allele. Therefore, potential targets for KASP marker development were chosen by filtering SNPs in this region to ensure that all 13 prickle-free genotypes were homozygous for the alternate allele and all four genotypes with prickles had one to four copies of the reference allele. Eight SNPs were chosen for Kompetitive Allele-Specific PCR (KASP) assay development at LGC Genomics (Beverly, MA, USA) based on proximity to the most significant SNP identified in GWAS. Each assay was designed to target a SNP with two forward primers and one common reverse primer (Supplementary Table 3). The two allele-specific forward primers targeting each were designed with a unique tail sequence labeled with universal FRET (fluorescence resonant energy transfer) cassettes, FAM or HEX dye.

A total of 626 diverse blackberry cultivars and breeding selections with known phenotypes were genotyped with the eight KASP markers to determine which marker was most predictive and broadly applicable across multiple blackberry breeding programs (Supplementary Table 4). The 626 genotypes used for KASP validation were sourced from the UADA program (n=497), the United States Department of Agriculture – Horticultural Crops Production and Genetic Improvement Research Unit (USDA-ARS-HCPGIRU, n=77), Hortifrut Genetics (HFG, n=40), and the United States Department of Agriculture – National Clonal Germplasm Repository (USDA-ARS-NCGR, n=12). One hundred and seventy-four of the 626 genotypes in the validation panel were previously included in the GWAS (158 cultivars and breeding selections from UADA, four USDA-ARS-HCPGIRU cultivars, and all 12 historical cultivars and germplasm accessions from USDA-ARS-NCGR). The new materials that were not included in the initial GWAS were comprised of 73 breeding selections from USDA-ARS-HCPGIRU, 40 breeding selections from HFG, 45 additional breeding selections from UADA, and 334 seedlings from two biparental populations segregating for prickles in the UADA breeding program.

KASP genotyping was performed with LGC’s genotyping service (LGC Genomics, Beverly, MA, USA). The KASP assay for each sample was conducted in a 4 µl reaction, which included 2 µl low ROX KASP master mix, 0.106 µl of primer mix (0.318 ll of each primer at final concentration), and 2 µl of 10–25 ng µl^-1^ genomic DNA. The PCR conditions were an initial denaturation step at 94 °C for 15 min, followed by 10 cycles of touch-down PCR with annealing temperatures decreasing from 68 °C to 60 °C dropping at the rate of 0.8 °C per cycle, followed by 30 cycles of denaturation at 94 °C for 20 s and primer annealing at 57 °C for 1 min. PCR fluorescent endpoint data were read using the LightCycler 480 Real-Time PCR System (Roche, Germany). The fluorescence signals measured from each sample were used to create a cluster plot in the R package ggplot2 (Wickham 2016).

### Analysis of Allele Dosage on Prickle Density

Two biparental populations were used to explore allele dosage at the prickle-free locus and its potential effect on prickle density. The first population (1034) resulted from a controlled cross of A-2447 (prickly) × APF-138T (prickle-free), and the second population (1525) resulted from a controlled cross of APF-195T (prickle-free) × Prime-Ark^®^ Horizon (prickly). Both A- 2447 and Prime-Ark^®^ Horizon were derived from crosses between prickly and prickle-free parents and presumed to be heterozygous, with two prickly alleles and two prickle-free alleles. Seedlings from each population were planted at 60 cm spacing in the field in the spring of 2019.

One hundred and forty-seven individuals from each population were evaluated for prickle density in 2019 and 2020. Prickle density was measured by collecting three canes from each plant at 5 cm above soil level and counting the number of prickles on the proximal 50 cm of each cane. The 334 plants from populations 1034 and 1525 were then genotyped with the eight KASP markers developed in the prickle-free locus as described above. A generalized linear model (GLM) analysis was carried out using the Poisson family function to test the effect of allele dosage of the SNP markers on prickles density. Finally, contrasts between allele dosages were obtained using the estimated marginal means with the *emmeans* R package (Lenth 2024).

## Results

### Association Analysis

A total of 374 blackberry genotypes were evaluated for the presence or absence of prickles and used in the association study. Thirty-eight genotypes (10.2%) had prickles, and the remaining 336 genotypes (89.8%) were prickle-free (Supplementary Table 1). The genotypic data resulting from CaptureSeq analysis used in this study are described in detail by Chizk et al. (2023). Briefly, 65,995 SNPs with an average read depth of 216 across all genotypes passed quality filtering and were used in association analysis. Over 99% of SNPs used in the association analysis were in annotated genic regions, and SNP density closely mirrored patterns of gene density across the *R. argutus* genome (Chizk et al. 2023). QQ-plots did not show evidence of systemic bias for the models evaluated in this study (Supplementary Fig. 1). A single locus associated with the prickle-free trait was detected containing 684 significant SNPs from 30484542 to 36088182 bp on chromosome Ra04 (Fig. 1, Supplementary Table 5). The peak SNP in the prickle-free locus was at 33,636,565 bp and had an LOD value of 75.28 under the simplex dominant gene model. The peak marker correctly predicted the phenotype (prickly vs. prickle-free) of 365 of 374 genotypes (97.6%) in the association analysis panel (Supplementary Table 1). The diploid germplasm accession ‘Burbank Thornless’ was the only prickle-free genotype that was heterozygous at position 33,636,565 bp. Eight UADA breeding selections (S217, S252, S259, S260, S301, S326, S327, and S331) were phenotyped as prickly but had four copies of the allele associated with prickle-free canes (“TTTT”).

**Fig. 1.**
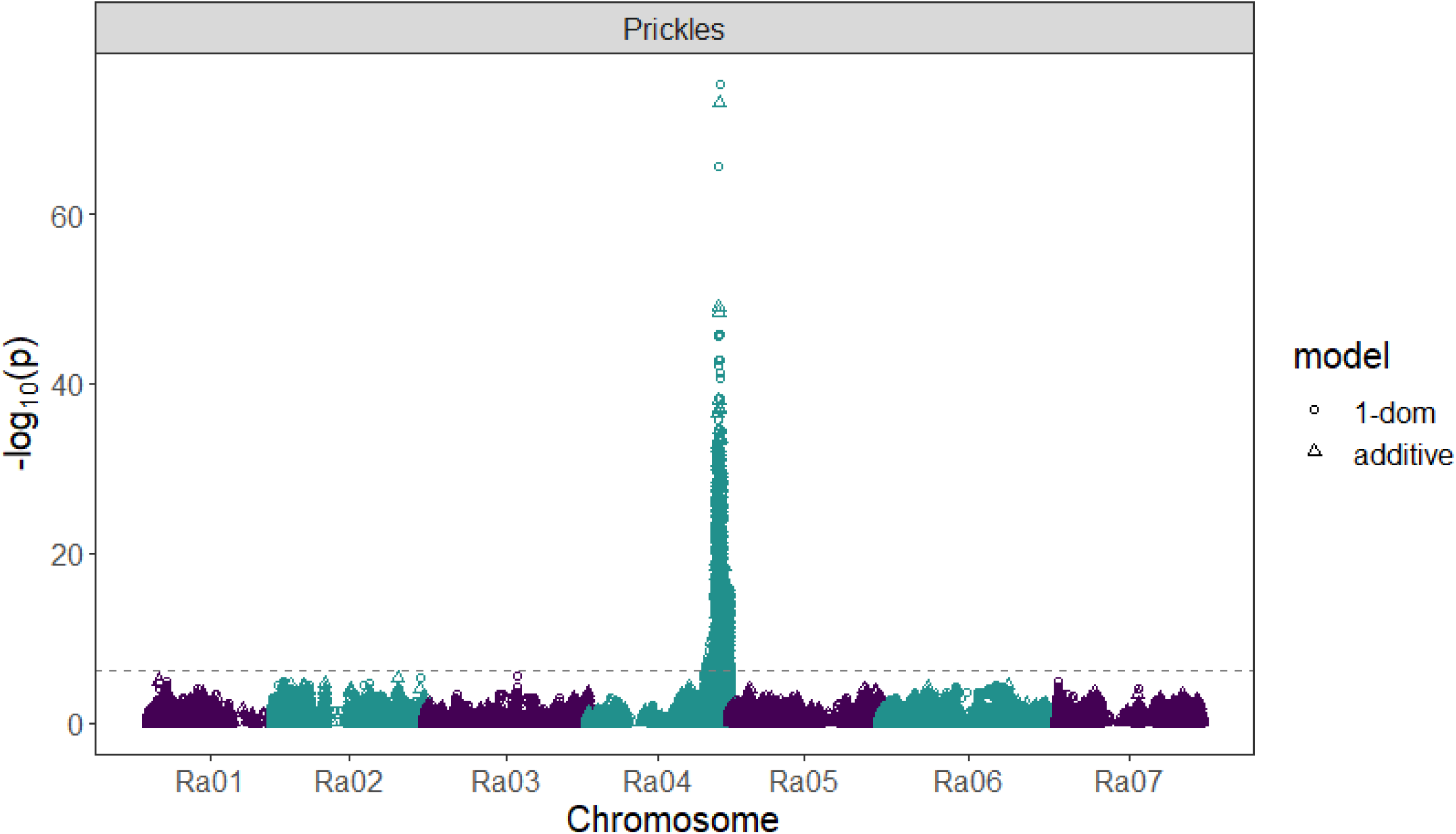
Manhattan plot showing association of 65,995 SNPs with blackberry prickles under additive and simplex-dominance models. The dashed line represents the lowest significance threshold used between tested models at α = 0.05 determined using the ‘M.eff’ method in GWASpoly.

### Possible Candidate Gene Identification

The rate of LD decay across chromosome Ra04 was highly variable, with a large block of extensive LD in the distal portion of the chromosome containing the prickle-free locus (Fig. 2). Therefore, possible candidate genes and transcription factors potentially associated with prickle development were investigated in a 5.6 Mb region extending from the first SNP significantly associated with the prickle-free locus at 30,484,542 to the distal end of chromosome Ra04. Five potential candidate genes with functional annotations related to epidermal, trichome, and/or prickle development were identified within the prickle-free locus (Table 1). A SQUAMOSA promoter-binding protein-like domain 6 (*SPL6*), Ra_g19233, was located at 32,260,895 to 32,263,859 bp, 1.3 Mb from the peak marker. A gene encoding a MYB domain protein 16 (*MYB16*), Ra_g19351, was located at 32,802,206 to 32,803,665 bp, 832 kb from the peak marker. An AGAMOUS-like MADS-box protein 30 (*AGL30*), Ra_g19365, was located at 32,877,483 to 32,880,709 bp, 755 kb from the peak marker. A homeobox-leucine zipper protein (*HOX3*), Ra_g19498, was located at 33,599,420 to 33,601,244 bp, 35 kb from the peak marker. Lastly, a trichome birefringence-like 27 protein (*TBL27*), Ra_g19762, was found at 34,904,416 to 34,905,885 bp, 1.26 Mb from the peak marker associated with prickles.

**Fig. 2.**
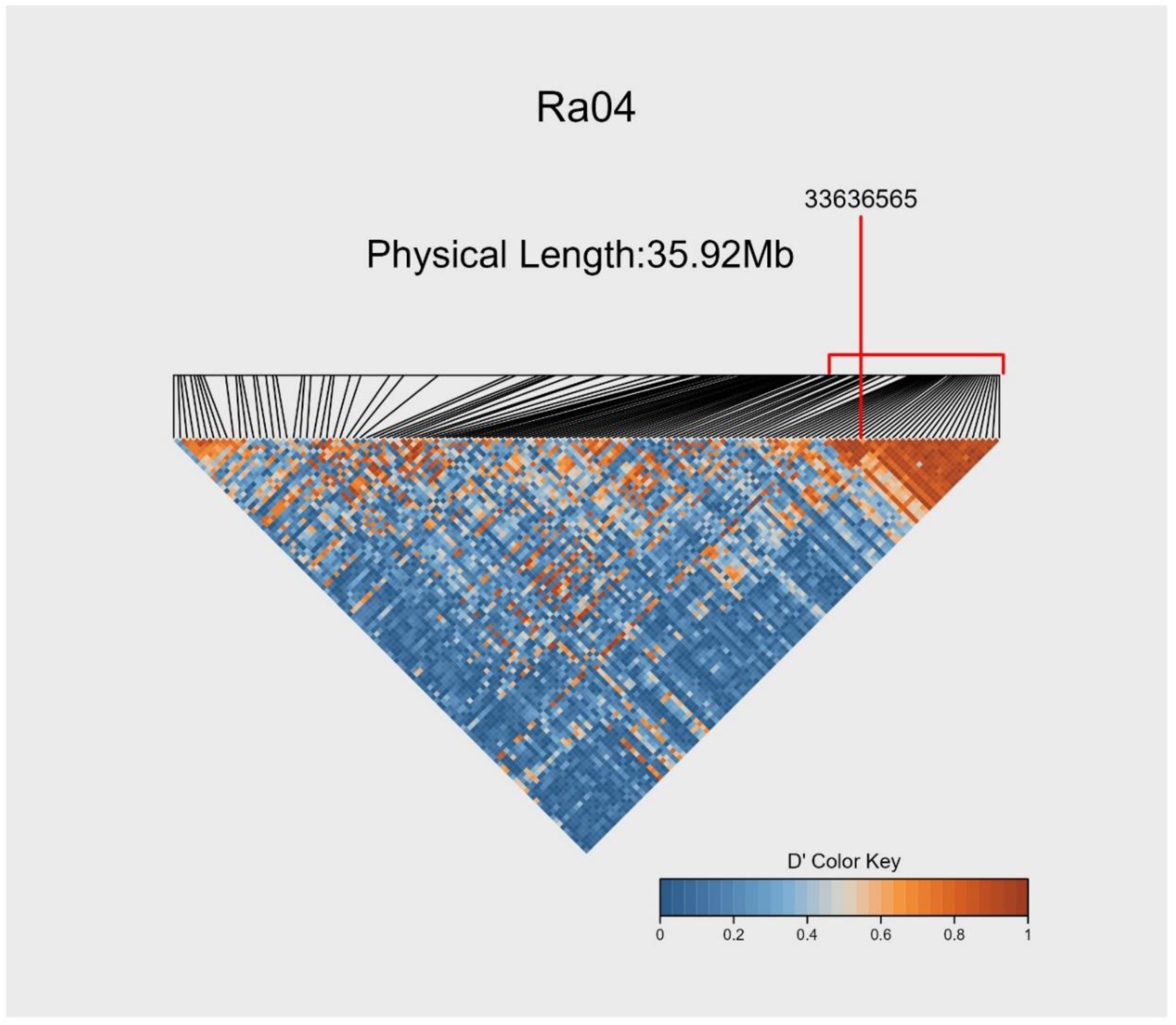
Heat map of absolute values of Lewontin’s D’ (Lewontin 1963) on blackberry chromosome Ra04 with SNPs subsampled to have a minimum spacing of 100 kb. The physical region containing 684 SNPs significantly associated with the prickle-free trait is indicated with a red horizontal line. The red vertical line indicates the physical position of the SNP most strongly associated with the prickle-free trait at 33,636,565 bp.

**Table 1.** Possible candidate genes for blackberry prickles located in a 5.6 Mb region associated with the prickle-free trait on chromosome Ra04

| Gene ID | Start bp | Stop bp | Functional annotation<sup>a</sup> | *Arabidopsis*<br>homolog |
| --- | --- | --- | --- | --- |
| Ra_g19233.t1 | 32260895 | 32263859 | SQUAMOSA promoter-binding-like protein 6 (SPL6) | At1g69170 |
| Ra_g19351.t1 | 32802206 | 32803665 | Myb domain protein 16 (MYB16) | At5g15310 |
| Ra_g19365.t1 | 32877483 | 32880709 | AGAMOUS-like MADS-box protein 30 (AGL30) | At2g03060 |
| Ra_g19498.t1 | 33599420 | 33601244 | homeobox-leucine zipper protein HOX3 (HD-ZIP IV) | At2g01430 |
| Ra_g19762.t1 | 34904416 | 34905885 | Trichome birefringence-like 27 | At1g70230 |

<sup>a</sup>The SwissProt and Araport11 databases were interrogated for descriptions of gene homologs and the *Arabidopsis* homolog identifier as described by Bruna et al. (2023).

### Whole Genome Sequencing

Whole genome sequencing of the seventeen tetraploid blackberry genotypes yielded between 92 and 123 million raw paired-end reads. After quality filtering and removing low-quality bases, an average of 106 million paired-end reads were produced per genotype (Supplementary Table 2). Sequence depth was 54X on average, ranging from 45X for S9 and Black Magic^TM^ to 62X for S17. The five potential candidate genes in the region of the prickle- free locus with functional annotations related to epidermal, trichome, and/or prickle development were surveyed for nonsynonymous mutations or structural variants that might cause the prickle-free trait. However, no such polymorphisms were found within the coding regions of these genes that were homozygous for the alternate allele in the 13 prickle-free genotypes and had one or more copies of the reference allele in all four thorny genotypes. A total of 1,479 SNPs and small indels were found in a region of 700 kb around the peak SNP associated with the prickle-free trait using whole genome sequencing data. Of these polymorphisms, 760 (51%) fit the expected pattern with all 13 prickle-free genotypes homozygous for the alternate allele and all four prickly genotypes with one to four copies of the reference allele. Based on these results, eight candidate SNPs at positions 33,572,441 bp, 33,573,461 bp, 33,581,279 bp, 33,582,533 bp, 33,583,532 bp, 33,600,544 bp, 33,600,857 bp, and 33,636,565 bp on chromosome Ra04 were selected for KASP development (Supplementary Table 3). While the CaptureSeq results predicted that all genotypes in the association panel were either homozygous (“TTTT”) or triplex (“TTTA”) for the SNP at 33,583,532 bp, WGS data showed that the prickly genotypes varied in their allele dosage. Black Magic^TM^ had only prickly allele (“TTTA”), Kiowa and Prime-Ark® 45 had two prickly alleles (“TTAA”), and Tupy had three prickly alleles (“TAAA”).

### KASP Validation

Of the eight KASP markers initially tested, two were found to be highly predictive in the validation panel (Fig. 3, Supplementary Table 4). These markers, which targeted SNPs at 33,572,441 and 33,636,565 bp on chromosome Ra04 were named *thorn1* and *thorn2,* respectively. The *thorn1* marker reaction failed for 22 of 626 genotypes in the validation panel. Of the 604 genotypes with successful marker reactions, the marker correctly predicted the phenotype of 586 (97%). Two seedlings from population 1034 (1034-056 and 1034-149), two seedlings from population 1,525 (1525-085 and 1525-086), and eight UADA breeding selections (S23, S57, S84, S132, S149, S156, S164, and S191) were scored as prickle-free but predicted to have one prickly allele by *thorn1*. Additionally, four USDA-ARS-HCPGIRU breeding selections were phenotyped as prickle-free but predicted to have one (OR17), three (OR33, OR35), or four (OR34) prickly alleles. Two breeding selections from the USDA-ARS-HCPGIRU program (OR05 and OR44) had all four copies of the prickle-free allele at *thorn1* but were phenotyped as prickly.

**Fig. 3.**
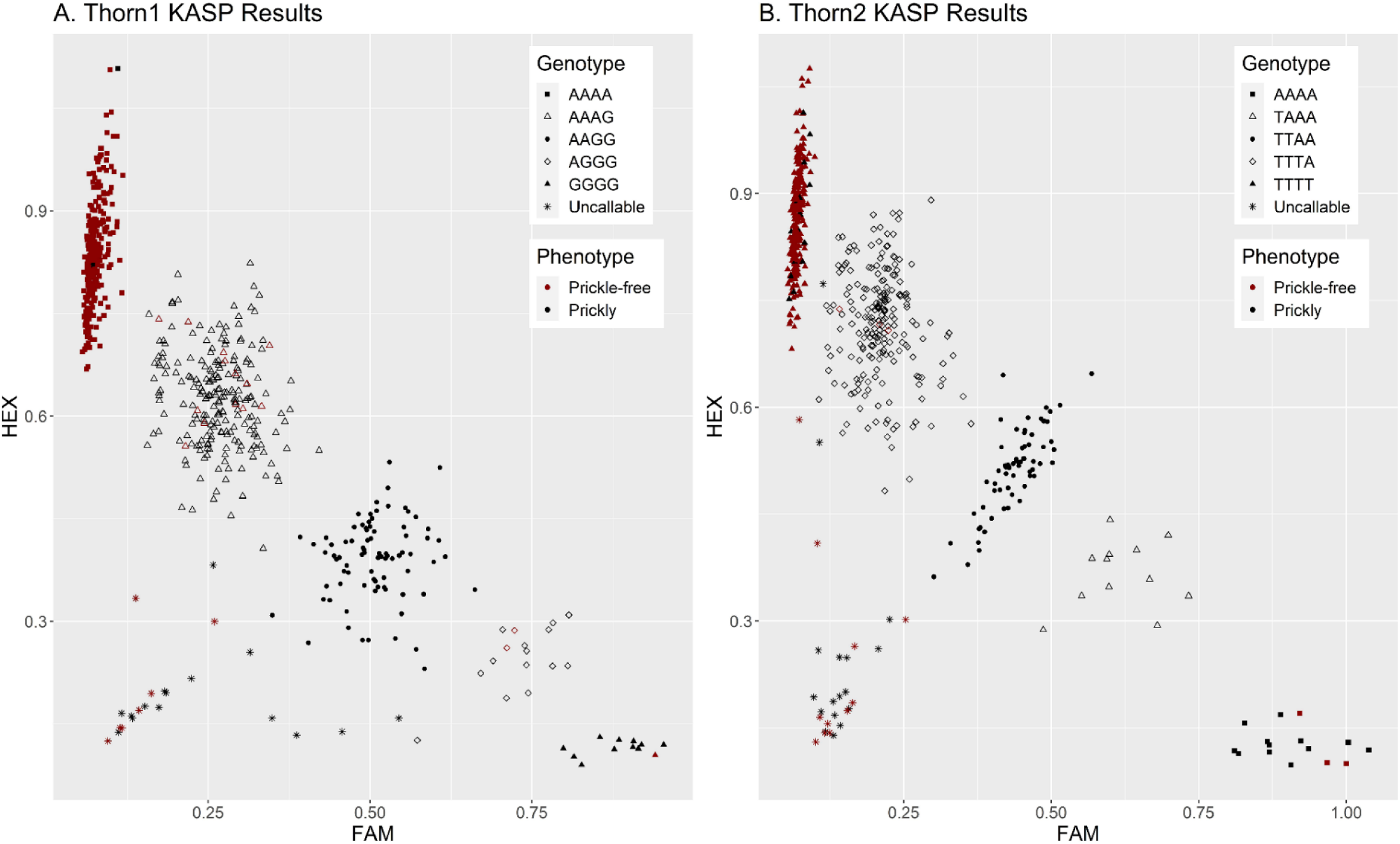
Scatter plot showing KASP markers results from a validation panel of 626 genotypes for *thorn1*, which targets a SNP at 33,572,441 bp on Ra04, and *thorn2*, which targets the peak SNP most strongly associated with the prickle-free trait in the GWAS at 33,636,565 bp on Ra04. Each cluster represents a different allele dosage class. Prickle-free genotypes are colored red, while prickly genotypes are colored black.

Twenty-eight of the 626 marker reactions failed for the *thorn2* marker. Of the 598 successful reactions, the *thorn2* marker correctly predicted the phenotype of 572 (96%) (Supplementary Table 4). One seedling from population 1034 (1034-149) and two seedlings from population 1525 (1525-085 and 1525-086) were scored as prickle-free but predicted to have one prickly allele by *thorn2*. The same four USDA-ARS-HCPGIRU breeding selections were phenotyped as prickle-free but predicted to have one prickly allele by *thorn1* and were also predicted to be prickly by *thorn2.* The prickle-free selection OR17 was predicted to have one prickly allele, while OR33, OR34, and OR35 were all predicted to have all four prickly alleles.

Nineteen genotypes in the validation panel had four prickle-free alleles for *thorn2* but were phenotyped as prickly. These genotypes included one seedling from population 1034 (1034-009), seven breeding selections from the UADA program (S217, S252, S259, S301, S326, S327, and S331), and eleven breeding selections from the USDA-ARS-HCPGIRU program (OR03, OR04, OR05, OR06, OR11, OR12, OR20, OR40, OR44, OR45, and OR46).

### Analysis of Marker Dosage

The effect of allele dosage for the *thorn1* and *thorn2* markers was evaluated in two biparental populations (1034 and 1525). The number of plants with prickly (n = 122) and prickle-free (n = 25) canes in population 1525 fit the expected 5:1 ratio for a cross between a duplex prickly parent and a prickle-free parent under random chromosome assortment (χ^2^ = 0.01, *P =* 0.91) (Table 2, Supplementary Table 6). For the 135 progeny in population 1525 with successful reactions for *thorn1*, the predicted genotype classes also fit the expected ratio of 1:4:1 (χ^2^ = 4.39, *P =* 0.11), with 19 “AAAA” genotypes, 101 “AAAG” genotypes, and 15 “AAGG” genotypes (Table 2). However, the results for marker *thorn2* in population 1525 were slightly distorted compared to the expected 1:4:1 ratio (χ^2^ = 6.19, *P =* 0.05). In this case, there was a slight excess of “TTTA” genotypes (n = 102) relative to “TTTT” (n = 19) and “TTAA” (n = 13) genotypes compared to the expected 1:4:1 ratio.

**Table 2.**
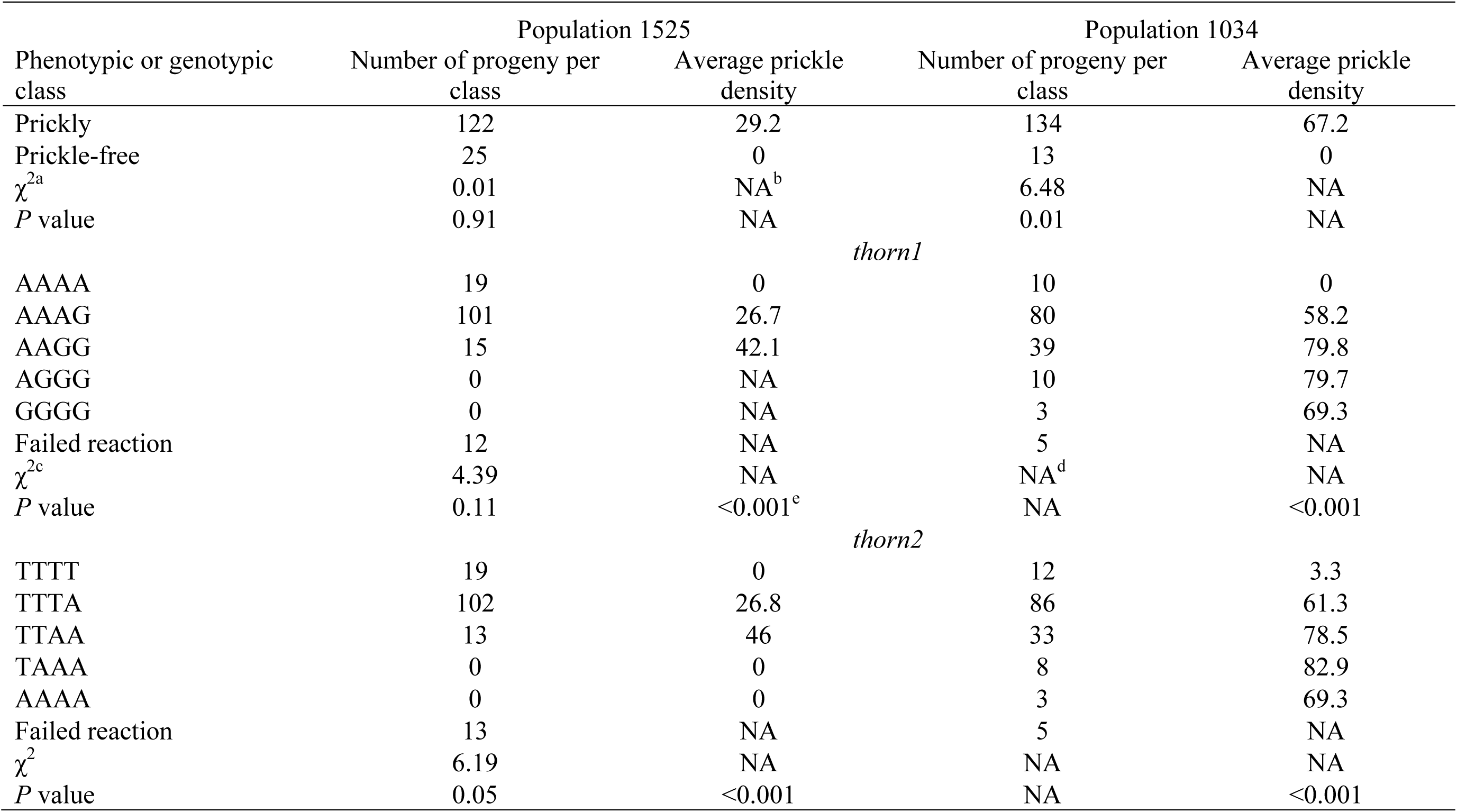
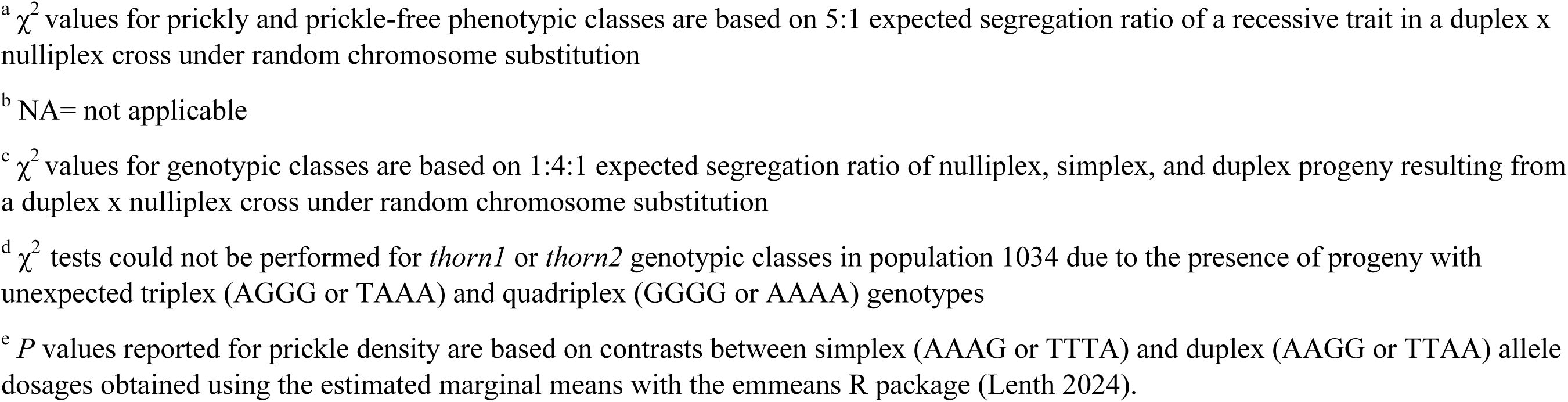
Summary of Chi Square (Χ^2^) tests for number of prickly and prickle-free progeny and genotype classes in populations 1525 and 1034 along with average prickle density for each population and class and *P* values for contrasts between simplex (AAAG or TTTA) and duplex (AAGG or TTAA) allele dosages obtained using the estimated marginal means.

| Phenotypic or genotypic class | Population 1525 |  |  | Population 1034 |  |
| --- | --- | --- | --- | --- | --- |
|  | Number of progeny per class | Average prickle density |  | Number of progeny per class | Average prickle density |
| Prickly | 122 | 29.2 |  | 134 | 67.2 |
| Prickle-free | 25 | 0 |  | 13 | 0 |
| $\chi^2$<sup>a</sup> | 0.01 | NA<sup>b</sup> | | 6.48 | NA |
| $P$ value | 0.91 | NA | | 0.01 | NA |
|  |  |  | *thorn1* |  |  |
| AAAA | 19 | 0 |  | 10 | 0 |
| AAAG | 101 | 26.7 |  | 80 | 58.2 |
| AAGG | 15 | 42.1 |  | 39 | 79.8 |
| AGGG | 0 | NA |  | 10 | 79.7 |
| GGGG | 0 | NA |  | 3 | 69.3 |
| Failed reaction | 12 | NA |  | 5 | NA |
| $\chi^2$<sup>c</sup> | 4.39 | NA | | NA<sup>d</sup> | NA |
| $P$ value | 0.11 | <0.001<sup>e</sup> | | NA | <0.001 |
|  |  |  | *thorn2* |  |  |
| TTTT | 19 | 0 |  | 12 | 3.3 |
| TTTA | 102 | 26.8 |  | 86 | 61.3 |
| TTAA | 13 | 46 |  | 33 | 78.5 |
| TAAA | 0 | 0 |  | 8 | 82.9 |
| AAAA | 0 | 0 |  | 3 | 69.3 |
| Failed reaction | 13 | NA |  | 5 | NA |
| $\chi^2$ | 6.19 | NA | | NA | NA |
| $P$ value | 0.05 | <0.001 | | NA | <0.001 |

<sup>a</sup> $\chi^2$ values for prickly and prickle-free phenotypic classes are based on 5:1 expected segregation ratio of a recessive trait in a duplex x nulliplex cross under random chromosome substitution
<sup>b</sup> NA= not applicable
<sup>c</sup> $\chi^2$ values for genotypic classes are based on 1:4:1 expected segregation ratio of nulliplex, simplex, and duplex progeny resulting from a duplex x nulliplex cross under random chromosome substitution
<sup>d</sup> $\chi^2$ tests could not be performed for *thorn1* or *thorn2* genotypic classes in population 1034 due to the presence of progeny with unexpected triplex (AGGG or TAAA) and quadriplex (GGGG or AAAA) genotypes
<sup>e</sup> *P* values reported for prickle density are based on contrasts between simplex (AAAG or TTAA) and duplex (AAGG or TTAA) allele dosages obtained using the estimated marginal means with the emmeans R package (Lenth 2024).

For population 1034, there was an excess of prickly (n = 134) progeny and fewer prickle-free progeny (n =13) relative to the expected 5:1 ratio (χ^2^ = 6.48, *P =* 0.01) (Table 2, Supplementary Table 6). Considering that this was a cross between a duplex prickly parent and a nulliplex prickle-free parent, we expected to find progeny with nulliplex, simplex, and duplex genotypes in a 1:4:1 ratio. However, it was not possible to test whether the segregation ratio in the progeny fit expected ratios because of the presence of unexpected genotype classes for both markers. There were 10 progenies with triplex genotypes (“AGGG”) and three progeny with quadriplex genotypes (“GGGG”) for *thorn1* and eight progeny with triplex genotypes (“TAAA”) and three progeny with quadriplex genotypes (“AAAA”) for *thorn2* (Table 2). The female parent of population 1034 was duplex for the prickly allele, so it is possible that these unexpected genotype classes in the progeny resulted from accidental self-pollination or contaminated pollen from wild blackberries. Because of the uncertainty of their pedigrees, progenies with three and four prickly alleles were excluded from subsequent pairwise comparisons of prickle density among different genotype classes in population 1034.

Significant differences in prickle density were observed between different allele dosages in both populations, and for both molecular markers (Table 2; Fig. 4). As expected, progenies with no prickly alleles (genotype “AAAA” for *thorn1* and “TTTT” for *thorn2*) had zero prickles, with the exception of seedling 1034-009 for marker *thorn2* (Supplementary Tables 4 and 6).

**Fig. 4.**
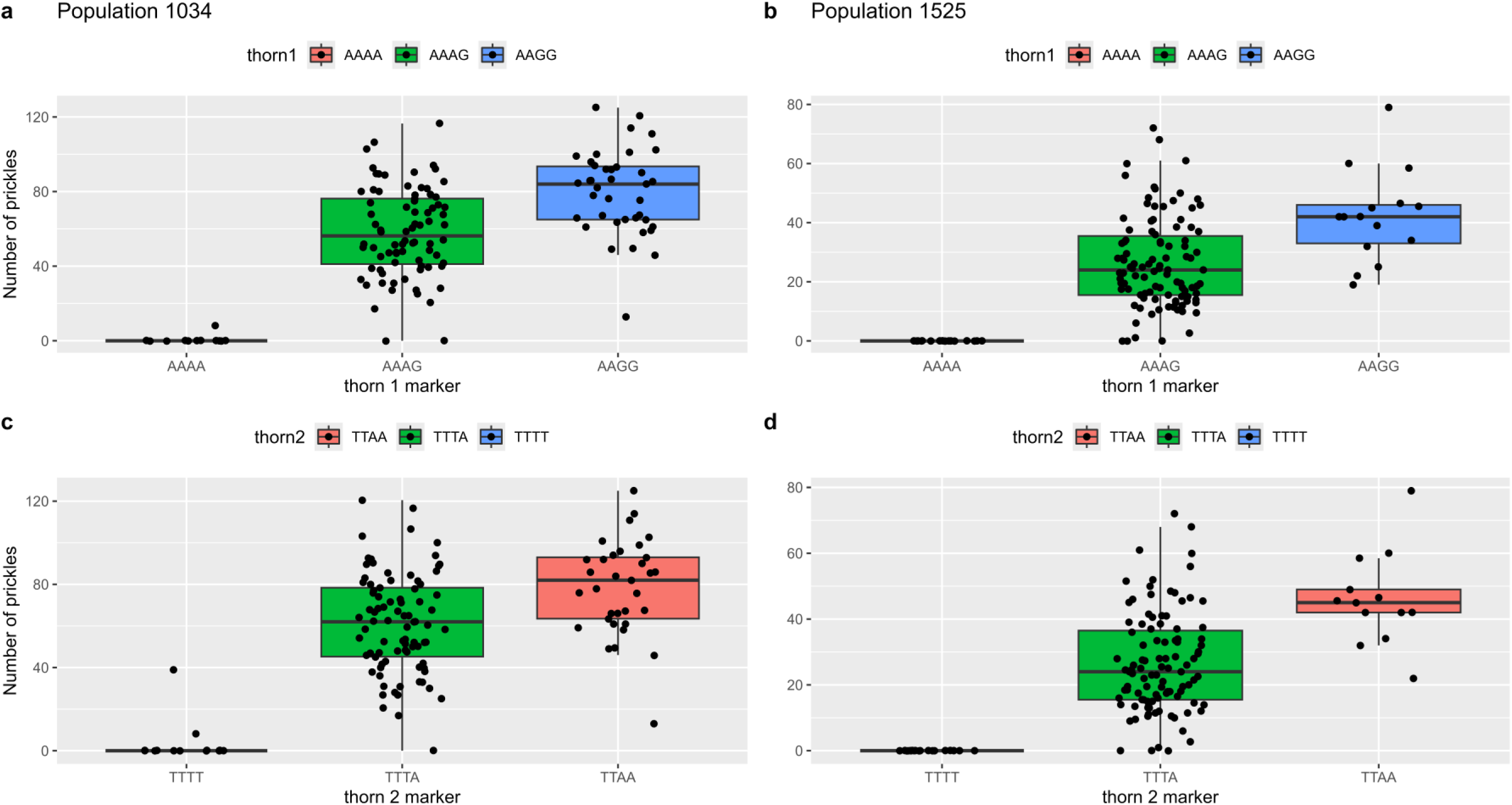
Effect of allele dosage on prickle density in two segregating biparental populations. Boxplots show the distribution of the number of prickles on a 50 cm cane segment for different allele dosages of *thorn1* (Ra04_33,572,441) and *thorn2* (Ra04_33,636,565) KASP markers in progeny from biparental population 1034 (panels A and C) and population 1525 (panels B and D). Each box represents the interquartile range (IQR), with the median indicated by the horizontal line.

Interestingly, in population 1034, progenies with two prickle alleles had a mean of 79.8±3.5 prickles (genotype “AAGG” for *thorn1*) and 78.5±4 prickles (genotype “TTAA” for *thorn2*), while progenies with only one prickle allele had a mean of 58.2±2.5 prickles (genotype “AAAG” for *thorn1*) and 61.3±2.4 prickles (genotype “TTTA” for *thorn2*). A similar pattern was observed in population 1525, where progenies with genotype “AAGG” for *thorn1* had a mean of 42.1±3.6 prickles, and those with genotype “TTAA” for *thorn2* had a mean of 46.0±4.0 prickles.

Progenies with genotype “AAAG” for *thorn1* and “TTTA” for *thorn2* showed means of 26.7±1.4 and 26.8±1.4 prickles, respectively (Table 2; Fig. 4). All pairwise comparisons indicated a significant effect of allele dosage on prickle density, with progeny with duplex genotypes having a higher density of prickles than simplex genotypes (Table 2).

## Discussion

A major locus associated with the ‘Merton Thornless’ source of the prickle-free trait in tetraploid blackberry was identified on chromosome Ra4 from 30.48 to 36.08 Mb in this study. A previous biparental mapping study in tetraploid blackberry conducted by Castro et al. (2013) placed the ‘Merton Thornless’ prickle-free locus in a large region on the distal portion of LG4.

The prickle-free locus was flanked by the SSR markers and RH_MEa0005aH07 and RH_MEa0011dG03b-265 at 4 and 21 cM in the map generated by Castro et al. (2013). The most tightly linked SSR marker, RH_MEa0005aH07, aligned between 34,851,955 and 34,852,561 bp on chromosome Ra04 of *R. argutus*. The prickle-free locus in diploid red raspberry was also placed in a 1.6 Mb (33.54–35.15 Mb) region on chromosome 4 of an unreleased red raspberry genome using an association analysis performed in a biparental population (Khadgi and Weber 2021). It is not possible to compare the exact physical position of the prickle-free locus identified in this study with previously mapped loci because the red raspberry genome used by Khadgi and Weber (2021) is not publicly available. However, orthologs of three of the possible candidate genes identified in this study, *MYB16*, *AGL30*, and *TBL27*, were recently proposed in red raspberry as possible candidates related to prickles based on their proximity to significant markers (Khadgi and Weber 2021). Overall, the placement of the prickle-free locus on chromosome Ra04 at 30.48-36.08 Mb corroborates previous mapping studies and suggests that the genetic control of prickle-free traits in red raspberry and ‘Merton Thornless’-derived tetraploid blackberries is similar.

### Peak Markers and Predictive KASP Assays

The peak SNP associated with prickles at 33,636,565 bp on chromosome Ra04 was strongly associated with the prickle-free trait; only nine out of 374 genotypes were incorrectly predicted. Of the nine genotypes inaccurately predicted by the SNP, only ‘Burbank Thornless’ had a prickle-free phenotype but was predicted to be heterozygous for the prickly allele. ‘Burbank Thornless’ (*R. ulmifolius inermis*, PI 554060), is a semi-erect diploid prickle-free blackberry accession. A prickle-free diploid *R. ulmifolius* plant was used as the original source of the prickle-free trait in ‘Merton Thornless’, but it is unclear if ‘Burbank Thornless’ is a clone of the *R. ulmifolius* plant used in the development of ‘Merton Thornless’. Additionally, eight prickly UADA breeding selections were predicted to have four copies of the prickle-free allele (S217, S252, S259, S260, S301, S326, S327, and S331). Intriguingly, all eight of these prickly breeding selections are closely related and were recently derived from a prickly wild Lebanese blackberry plant that was first used in crosses at UADA in 2009. S217 is a parent of S252, S259, S260, and S301 and a grandparent of S326, S327, and S331.

KASP markers targeting the peak SNP from GWAS at 33,636,565 bp on chromosome Ra04 (*thorn2*) and an additional SNP at 33,572,441 on chromosome Ra04 (*thorn1*) were highly predictive of the prickle-free trait in a validation panel composed of 626 cultivars, selections, and seedlings from the USDA-ARS-NCGR and the UADA, USDA-ARS-HCPGIRU, and HFG breeding programs (Fig. 3, Supplementary Table 4). *Thorn1* and *thorn2* markers correctly predicted the phenotype of 97% and 96%, respectively, of genotypes in the validation panel.

Interestingly, the results of the *thorn2* marker, which was designed to target the peak SNP discovered in GWAS at 33,636,565 bp, did not match CaptureSeq results for ‘Burbank Thornless.’ ‘Burbank Thornless’ was predicted to be heterozygous for the prickly allele at 33,636.565 bp in CaptureSeq, yet it grouped with the homozygous prickle-free group in the *thorn2* KASP analysis. Seven of the eight UADA breeding selections with prickly phenotypes that were predicted to be prickle-free by the peak SNP at 33,636,565 bp in CaptureSeq were also included in the KASP validation panel. All seven prickly genotypes derived from crosses with a prickly wild Lebanese blackberry plant (S217, S252, S259, S301, S326, S327, and S331) were also predicted to be prickle-free by *thorn2.* However, these seven selections were all correctly predicted to have either one or two prickly alleles by the *thorn1* marker, suggesting possible recombination between these SNPs. It is unclear if the genetic mechanism controlling the prickle-free trait is different from the rest of the GWAS panel in ‘Burbank Thornless’ and the Lebanese-derived germplasm or if recombination between the candidate gene controlling prickle development and the peak SNP at 33,636,565 bp limit the predictive ability of the *thorn2* in these genotypes. While the *thorn1* marker correctly predicted the phenotype of the seven prickly

Lebanese-derived UADA breeding selections, it predicted that eight prickle-free UADA breeding selections (S23, S57, S84, S132, S149, S156, S164, and S191) had one or more prickly allele. Both CaptureSeq genotyping results at 33,636,565 bp and the *thorn2* marker predicted these eight selections had four prickle-free alleles, indicating likely recombination between *thorn1* and the candidate gene for the prickle-free trait.

Eleven breeding selections from the USDA-ARS-HCPGIRU program (OR03, OR04, OR05, OR06, OR11, OR12, OR20, OR40, OR44, OR45, and OR46) were phenotyped as prickly but predicted as prickle-free by *thorn2*. Of these eleven prickly USDA-ARS-HCPGIRU genotypes, only two (OR05 and OR44) were also predicted to be prickle-free for *thorn1*. OR05 and OR44 are both tetraploid semierect selections with interesting pedigrees. OR05 has hexaploid *R. ursinus* and *R. caucasicus* in its pedigree record, and one grandparent of OR44 is a wild *R. insularis* accession from Northern Europe. Thus, it is possible that additional alleles or loci are impacting the prickly phenotype observed in these two selections. Four other breeding selections from the USDA-ARS-HCPGIRU (OR17, OR33, OR34, and OR35) were phenotyped as prickle-free but predicted to have one or more prickly alleles by both *thorn1* and *thorn2* KASP markers. OR33, OR34, and OR35 are all trailing blackberries with *R. ursinus* genetics and ‘Lincoln Logan’ in their pedigrees. ‘Lincoln Logan’ is a non-chimeral plant produced from tissue culture of the thornless chimera ‘Loganberry’ (Hall et al. 1986). Unlike ‘Merton Thornless’, the thornless allele from ‘Lincoln Logan’ is dominant (Hall and Stephens 1999). Therefore, it is likely that the prickle-free phenotype in these three selections is conferred by the ‘Lincoln Logan’ source and controlled by a different locus. On the other hand, OR17 is an erect tetraploid blackberry genotype without any exotic species in its recent pedigree. This genotype was predicted to have one thornless allele by both *thorn1* and *thorn2* markers and might be an interesting selection to use in fine mapping of the prickle-free locus.

### Effect of Allele Dosage on Thorn Density

The density of prickles varies widely among prickly fresh market blackberry germplasm (Scott et al. 1957; Pavlis and Moore 1981). Pavlis and Moore (1981) tested the hypothesis that cane prickle density was impacted by additive gene dosage at the major prickle-free locus inherited from ‘Merton Thornless’ in fourteen segregating and non-segregating seedling populations with representing quadriplex, triplex, duplex, simplex, nulliplex genotypes. Because the prickle-free locus had not been mapped at the time and no molecular markers were available to predict allele dosage in individual seedlings, Pavlis and Moore compared the mean, variance, and distribution curve of prickle density in each population. They found no significant difference in mean thorn density of populations expected to have all quadriplex prickly progeny from those expected to have all duplex prickly progeny. Furthermore, the variances and distribution curves of prickle density in segregating families expected to contain quadriplex, triplex, and duplex progeny were similar to those from nonsegregating families generated from crosses between two quadriplex parents or nulliplex and quadriplex parents. Based on these results, Pavlis and Moore (1981) concluded that prickle density was likely controlled by several modifying genes.

Similarly, Rawandoozi et al. (2024) used a pedigree-based QTL approach and identified 12 loci on four chromosomes impacting stem and leaf rachis prickle density in two multiparental diploid rose populations.

Our results indicate that allele dosage at the prickle-free locus in tetraploid blackberries has a significant impact on prickle density. Genotypes with two prickly alleles (“AAGG” for *thorn1* and “TTAA” for *thorn2*) had significantly higher prickle density than genotypes with only one prickly allele (“AAAG” for *thorn1* and “TTTA” for *thorn2*) in both populations 1034 and 1525. It is likely that Pavlis and Moore (1981) were unable to observe this effect of allele dosage in their populations because they were unable to predict the genotype of specific progeny and had to compare population means and variances to make inferences about the effect of allele dosage. It is also unclear whether genotypes with three or four prickly alleles would have higher prickle density than those with two prickly alleles. The association panel used in this study is poorly suited to study the effect of higher allele dosages on prickle density; only 38 of 336 genotypes were prickly and most of those had two or fewer prickly alleles. However, this would be an interesting trait to study in a different panel with more prickly germplasm. It should also be noted that there was a great deal of variation within each genotype class and that the progeny of population 1034 had over two times the prickle density of population 1535 on average despite both populations being derived from duplex (prickly) by nulliplex (prickle-free) parental combinations. Thus, our results suggest that other modifying loci likely also impact prickle density in tetraploid blackberries.

### Potential Candidate Genes for Prickle-Free Blackberries

An extensive LD block was identified around the prickle-free locus in the distal portion of chromosome Ra04. Large LD blocks like these have been associated with important domestication genes and corresponding selective sweeps in other crops (Kim and Stephan 2002). Indeed, the development of cultivars with prickle-free canes has been a major focus of most fresh-market blackberry breeding programs, and the selection intensity for this trait has been very high (Clark et al. 2007). The lack of recombination and reduced genetic diversity around the prickle-free locus may have negative implications for breeders. Moore (1984) found a strong linkage between prickle-free plants and trailing growth habit, acid fruit, and late harvest season.

Breaking such deleterious linkages may be more difficult if there is reduced recombination or overall allelic diversity in the distal portion of chromosome Ra04 in elite fresh-market blackberry germplasm. The extensive LD block around the prickle-free locus also complicates the search for candidate genes. For this reason, we surveyed a 5.6 Mb region from 30.48 Mb to the distal end of chromosome Ra04 for potential candidate genes with function annotations related to trichome or prickle development.

Prickles have been hypothesized to be modified trichomes (Kellogg et al. 2011; Khadgi and Weber 2020), although this is still unconfirmed, and alternative hypotheses have been proposed (Zhou et al. 2021). A relationship between stomata, trichomes, and prickle development has been proposed previously (Chalvin et al. 2020; Torii 2021). Khadgi and Weber (2021) proposed three possible candidate genes with functions related to trichome development near significant SNPs in the prickle-free locus in red raspberry: a *MYB16*-like transcription factor, the AGAMOUS-like MADS-box gene *AGL30*, and trichome birefringence-like 2 (*TBL2*). Two of the three candidates proposed in red raspberry (*MYB16* and *AGL30*) were located within the prickle-free locus in tetraploid blackberry. A homolog of *TBL2*, trichome birefringence 27 (*TBL27*) was found just outside the prickle-free locus and 1.27 Mb from the peak SNP associated with prickles in tetraploid blackberry. Two additional genes located on Ra04 between 30.48-34.31 Mb with functions related to trichome development, the SQUAMOSA promoter-binding-like protein 6 (*SPL6*) and the homeobox domain leucine zipper HOX3 (*HOX3*), are also proposed as possible candidate genes for the prickle-free trait.

The myleoblastosis (*MYB*) protein-coding genes belong to the *R2R3-MYB* subfamily of transcription factors (Chalvin et al. 2020). This family of proteins involves many functions related to stomatal regulation, trichome formation, and cuticle and wax formation (Baumann et al. 2007; Oshima and Mitsuda 2013; Chalvin et al. 2020; Yang et al. 2022). Two *R2R3-MYB* genes, *MYB16* and *MYB88*, were found within the QTL for blackberry prickles. *MYB16* regulates cuticle formation in reproductive organs and trichomes (Oshima and Mitsuda 2013; Oshima et al. 2013). Downregulation of *MYB16* is associated with reduced production of glandular trichomes, which could inhibit prickle development if prickles are indeed modified trichomes or share a related metabolic pathway as hypothesized (Baumann et al. 2007). In a transcriptome analysis, Khadgi and Weber (2020) found that MYB16 was significantly downregulated in the epidermis of prickle-free plants relative to prickly raspberry plants, making this an especially interesting candidate for the prickle-free trait. A gene encoding a MYB88 transcription factor protein (*MYB88*) (Ra_g19414) was also located at 33,143,949 to 33,147,906 bp within the QTL and 488 kb from the peak marker. *MYB88* has been described to function in stomata formation, and while not a likely candidate for prickles, this gene is interesting because of the proposed associations that exist in the origins of stomata, trichomes, and prickles (Torii 2021).

The possible candidate gene located nearest the peak SNP was a homeobox domain leucine zipper HOX3 (*HOX3*). *HOX3* is classified as an HD-ZIP IV, which has known functions in glandular trichome initiation and could be involved in R2R3-MYB/HD-ZIP IV complexes that differentiate trichome function (Chalvin et al. 2020). An established relationship exists between HOX3 and trichome development. In cotton (*Gossypium*), a *HOX3* transcription factor that is associated with trichome elongation has been identified (Shan et al. 2014).

*AGL30* is a transcription factor in the MADS-box family. Khadgi and Weber (2020) found that five MADS-box transcription factors were differentially expressed between prickle-free and prickly plants, and *AGL30* specifically was downregulated in the prickle-free epidermis of *Solanum viarum* (Pandey et al. 2018). Another little-studied but promising potential candidate gene for the prickle-free trait in blackberry was *TBL27*, a member of the trichome birefringence (*TBR*) gene family, which was located 1.27 Mb from the peak prickle-free marker. *TBR* functions in the formation and deposition of cellulose on the secondary cell wall (Potikha and Delmer 1995; Bischoff et al. 2010). Mutant *tbr Arabidopsis* plants were unable to generate secondary wall cellulose in trichomes found on leaves and stems (Potihka and Delmer 1995).

The last possible candidate gene for the prickle-free trait, *SPL6,* has been expressed at various degrees in transgenic *Arabidopsis* and tobacco (*Nicotiana tabacum*) plants. Mutants overexpressing the *SPL6* transcription factor had high trichome concentrations on leaf margins in contrast to the wild types, which had smooth rosette leaf margins (Ma et al. 2019). This is interesting, given that prickly tetraploid blackberries have trichomes on cotyledon leaf margins, while prickle-free blackberries have smooth cotyledon leaf margins. In maize (*Zea mays* L.), triple-knockout mutants of without functioning *ZmSPL10*, *ZmSPL14,* and *ZmSPL26* genes completely lacked trichomes but had higher densities of stomata, further confirming the existence of a common pathways for formation of stomata and non-stomata epidermal tissues (Kong et al. 2021).

All the identified gene candidates for the ‘Merton Thornless’ prickle-free trait in tetraploid blackberry have designated roles in stomata formation and trichome development and differentiation. Interestingly, whole genome resequencing data of four prickly and 13 prickle-free tetraploid blackberry genotypes did not reveal any nonsynonymous mutations or structural variants within these potential candidate genes that could be associated with the prickle-free trait. Transcriptome analysis specific to tetraploid blackberry should be performed to determine if these possible candidate genes are expressed at different levels in prickly and prickle-free genotypes.

## Conclusions

A single locus controlling prickle-free canes derived from ‘Merton Thornless’ was identified in tetraploid blackberry in this research. The prickle-free locus was located on chromosome Ra04 from 30.48 to 36.04 Mb in an extensive LD block. The peak SNP at 33.64 Mb correctly predicted the phenotype of all but nine of 374 cultivars and selections in the association panel. The location of the prickle-free locus is corroborated by other studies conducted in biparental raspberry and blackberry populations (Castro et al. 2013; Khadgi and Weber 2021). Five possible candidate genes and transcription factors with functional annotations related to epidermal, trichome, or prickle development were identified in the prickle-free locus, but no nonsynonymous polymorphisms associated with prickle-free canes were identified within these potential candidates using whole genome resequencing of 4 prickly and 13 prickle-free blackberries. While the causal gene controlling the prickle-free trait remains unknown, two useful KASP markers that are highly predictive of the presence/absence of prickles in diverse blackberry germplasm were developed in this study. Implementing marker-assisted selection for prickle-free canes can help breeders plan crosses, identify rare recombinants in the prickle-free locus for fine mapping, and cull prickly seedlings from populations before planting

## Data Availability

The whole genome resequencing data used in this study are publicly available at NCBI (accession number PRJNA1002337). The Capture-Seq derived SNP data used in GWAS are available in the Genome Database for Rosaceae repository (accession number tfGDR1069). All other data used in this study are presented in this manuscript and its supplementary files.

## Acknowledgments

The authors would like to thank Jackie Lee and the entire staff of the University of Arkansas System Division of Agriculture (UADA) Fruit Research Station for their hard work managing the plants used in this study. We also thank Arkansas Fruit Breeding graduate students Carly Godwin and Kenneth Buck for assisting with phenotyping and collaborating in the broader blackberry GWAS effort. We would like to thank Addison Eggert, Anderson Silva, and the RAPiD genomics team for their assistance and collaboration in CaptureSeq probe design, sequencing, and bioinformatics and LGC genomics for their work on KASP primer design.

## Author Contribution Statement

CAJ collected phenotype data, performed GWAS analysis, and drafted the manuscript in collaboration with MW. MW wrote the grant to support this research, supervised students and staff working on this project, and provided overall conceptual guidance. RA and HA prepared libraries for WGS and conducted bioinformatics to assist with Capture-Seq probe design. TMC performed tetraploid SNP calling, computed the Q matrix used in association mapping, and prepared Figure 2. AS analyzed WGS data, selected targets for KASP primer design, and analyzed the dosage effect on prickle density. LN performed DNA extractions and quantification. JRC developed most of the advanced selections and cultivars used in the study. ET and MH provided additional blackberry germplasm for KASP marker development and shared phenotype data. All authors contributed to the article and approved the submitted version.

---

**Supplementary Fig. 1.**
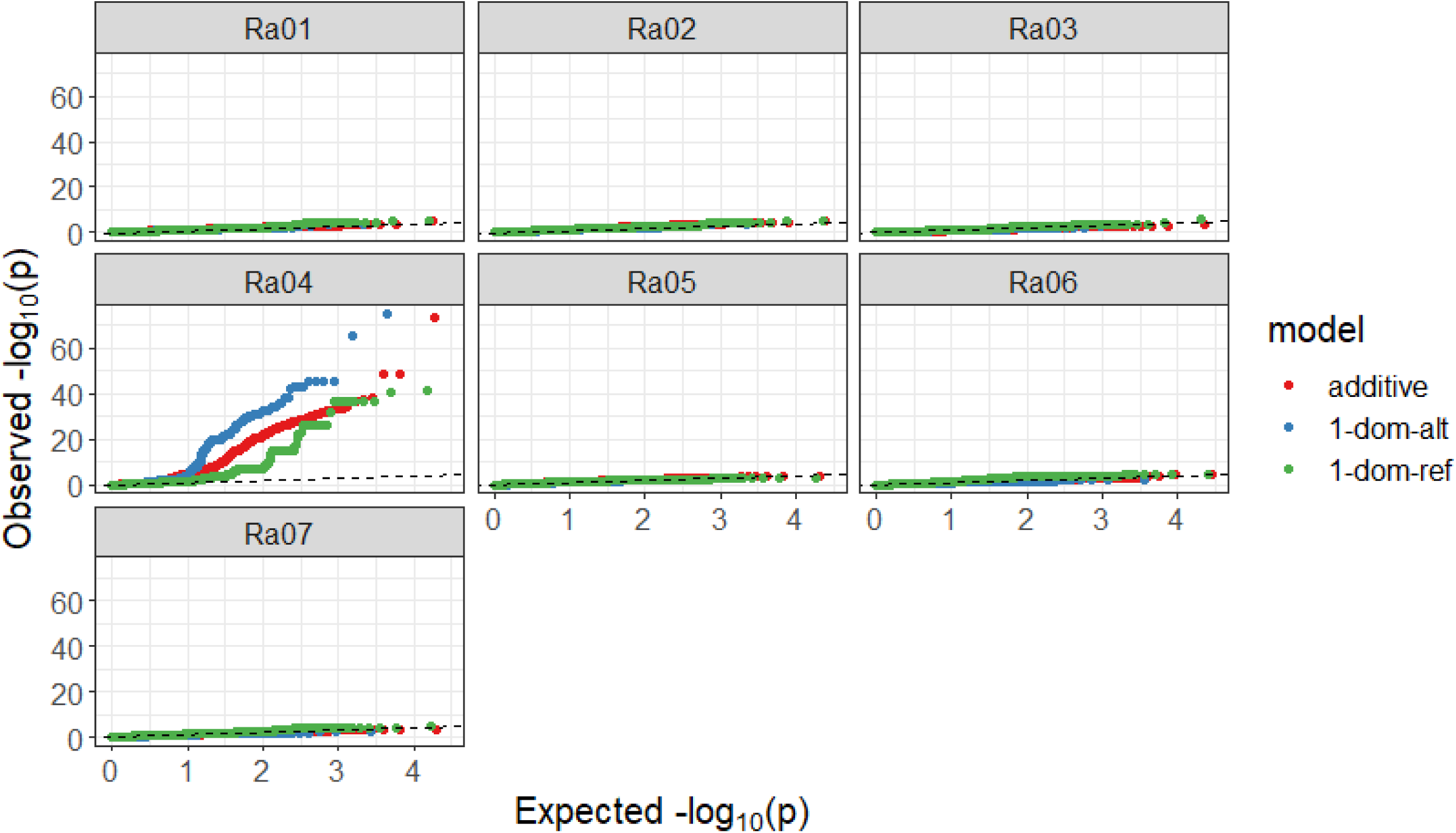
QQ-plots for prickles showing the observed versus expected distribution of p-values for markers on each *R. argutus* chromosome. Deviation from the fitted dashed line represents rejection of the null hypothesis that markers are not associated with traits.

